# Commentary on Pang et al. (2023) *Nature*

**DOI:** 10.1101/2023.07.20.549785

**Authors:** Joshua Faskowitz, Daniel Moyer, Daniel A. Handwerker, Javier Gonzalez-Castillo, Peter A. Bandettini, Saad Jbabdi, Richard Betzel

## Abstract

Pang et al. (2023) present novel analyses demonstrating that brain dynamics can be understood as resulting from the excitation of geometric modes, derived from the shape of the brain. Notably, they demonstrate that linear combinations of geometric modes can reconstruct patterns of fMRI data more accurately, and with fewer dimensions, than comparable connectivity-derived modes. Equipped with these results, and underpinned by neural field theory, the authors contend that the geometry of the cortical surface provides a more parsimonious explanation of brain activity than structural brain connectivity. This claim runs counter to prevailing theories of information flow in the brain, which emphasize the role of long-distance axonal projections and fasciculated white matter in relaying signals between cortical regions (Honey et al. 2009; Deco et al. 2011; Seguin et al., 2023). While we acknowledge that cortical geometry plays an important role in shaping human brain function, we feel that the presented work falls short of establishing that the brain’s geometry is “a more fundamental constraint on dynamics than complex interregional connectivity” (Pang et al. 2023). Here, we provide 1) a brief critique of the paper’s framing and 2) evidence showing that their methodology lacks specificity to the brain’s orientation and shape. Ultimately, we recognize that the geometric mode approach is a powerful representational framework for brain dynamics analysis, but we also believe that there are key caveats to consider alongside the claims made in the manuscript.

---

The claim made by Pang et al. rests largely on a comparison between brain shape and structural connectivity, in which modes derived from cortical surface geometry can more succinctly reconstruct functional brain maps than analogous connectivity modes. With these results in hand, the authors make claims that can be perceived as winner-takes-all, such as “if we prioritize spatial and physical constraints on brain anatomy, we only need to consider the shape of the brain, and not its full array of topologically complex axonal interconnectivity, to understand spatially patterned activity” and “while our findings cannot rule out a role for complex interregional connectivity they do indicate that such connectivity is not necessary for the emergence of these macroscale dynamics”. These claims raise the question: If cortical geometry shapes brain activity, what is the role of long-range structural connectivity?

A reconstruction of activation maps (Figs. 1 and 2 in Pang et al.) predicated on geometry must be reconciled with a century’s worth of observations wherein direct insults to white-matter pathways leave surface geometry intact but nonetheless result in acute changes in function, behavior, and cognition (Catani & ffytche, 2005; Filley & Fields, 2016). For example, how can the authors conciliate their framework with observations of acute functional changes following a callosotomy (O’Reilly et al. 2013), or distributed alterations to cortical activity following targeted pharmacogenetic disconnections of deep subcortical structures (Grayson et al. 2016)? Such questions illustrate some of the limitations of an account of brain activity that does not consider complex interregional white matter connectivity.

**Figure 1.**
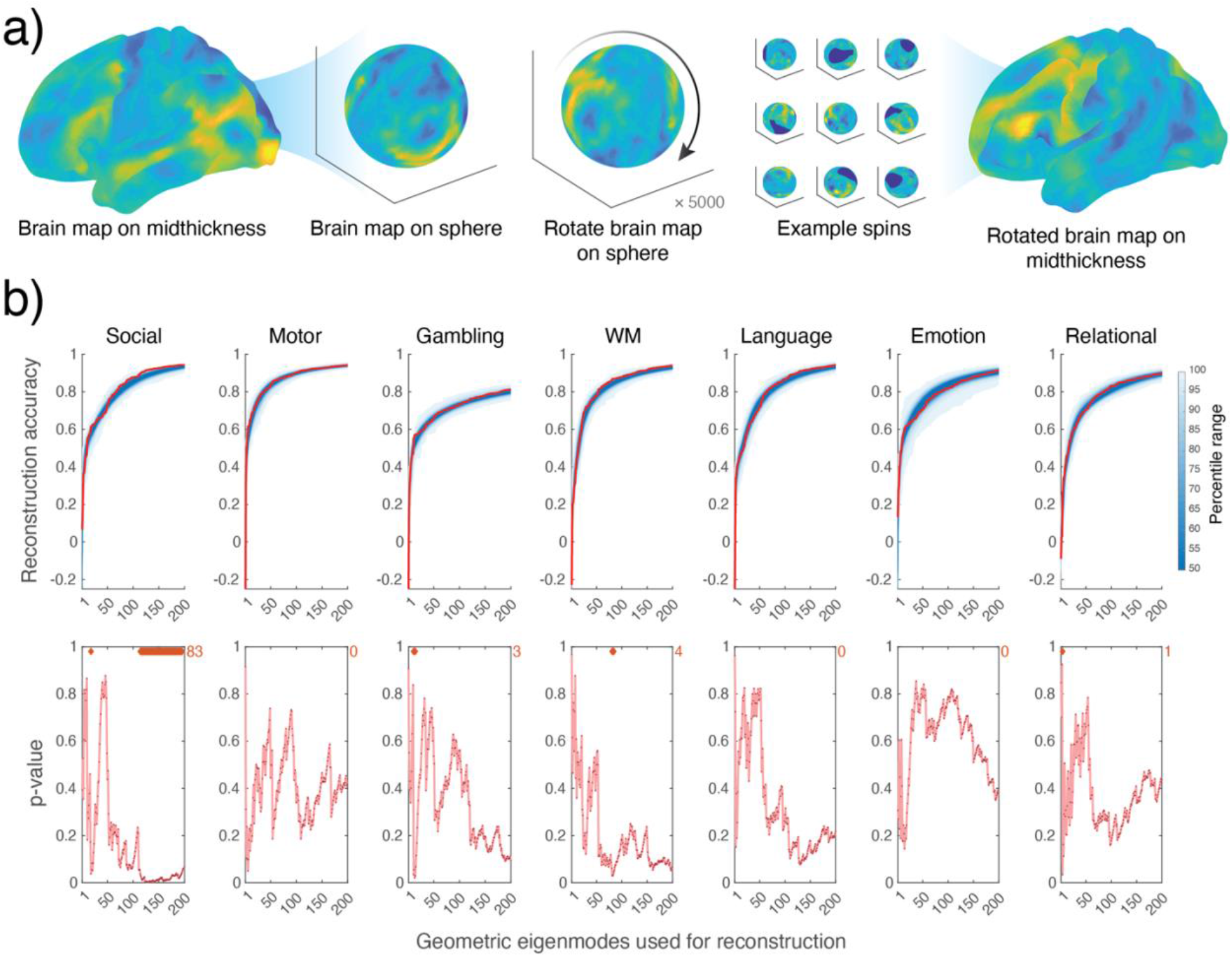
Using surface-based geometric eigenmodes to explain randomized contrast maps. (a) Schematic illustrating the procedure for generating randomized maps; task contrast maps are initially represented on midthickness surface, projected onto a sphere, which is randomly rotated before projecting the map back onto the midthickness surface. The resulting randomized map preserves the spatial statistics of the original map, but varies their locations. (b) We then use the geometric modes described by Pang et al. (2023) to explain the randomized maps for each of the seven task contrasts. The top row corresponds to reconstruction accuracy, which is the product-moment correlation between the empirical contrast data and the reconstructed contrast data with increasing numbers of eigenmodes. The red line indicates the reconstruction accuracy of the non-randomized data, as presented in Pang et al. (2023), whereas the blue shading indicates the percentile interval of the randomized data. The bottom row corresponds to p-value, the proportion of times that the empirical reconstruction accuracy exceeded the randomized reconstruction accuracy magnitudes, associated with each amount of modes; orange diamonds and counts indicate instances where p<0.05 (uncorrected for multiple comparisons).

**Figure 2.**
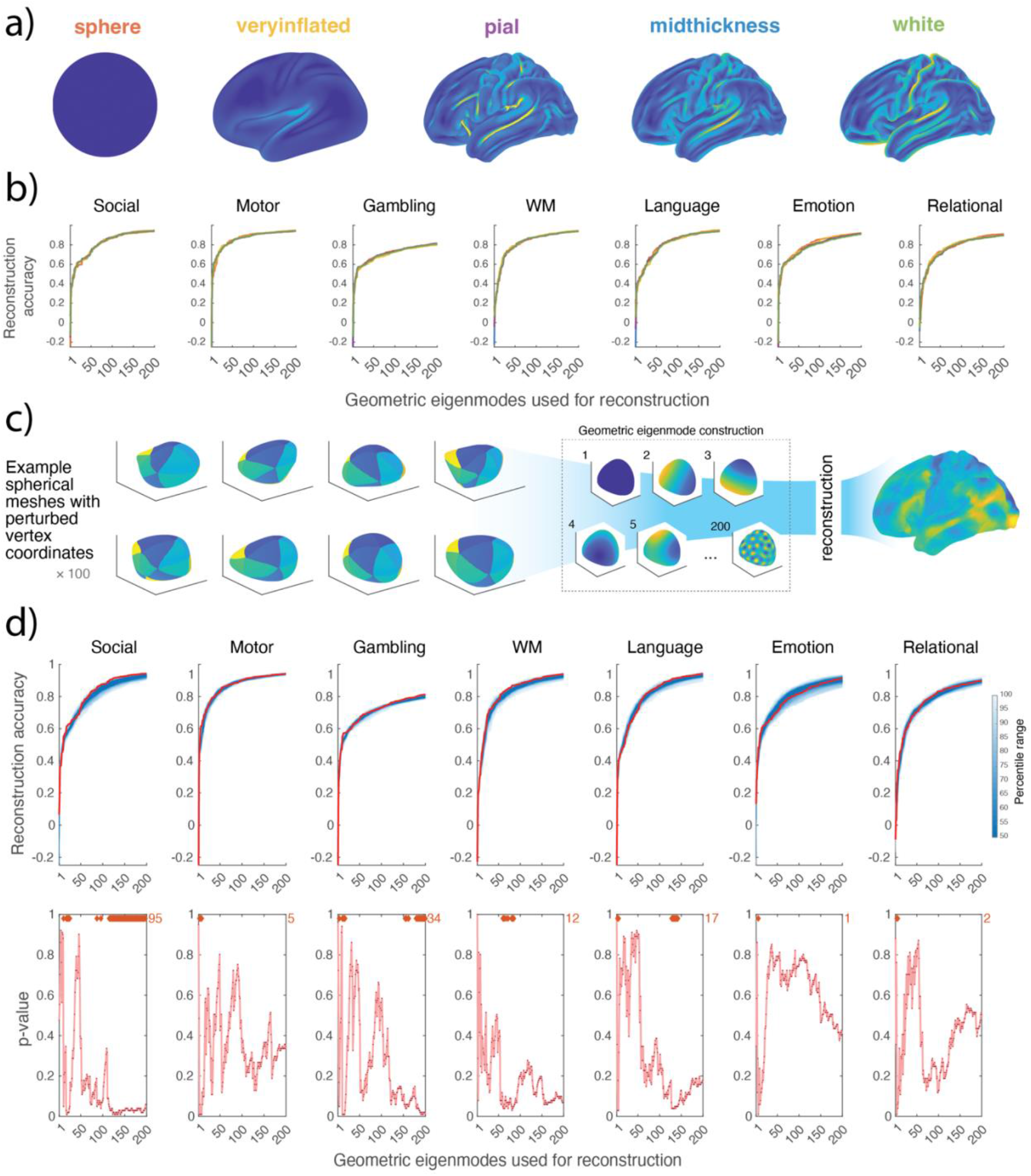
Evaluating the reconstruction accuracy of geometric eigenmodes derived from other surfaces. (a) Visualization of 32k surface meshes made available by the Human Connectome Project and which are commonly used in fMRI applications; surfaces are colored according to the absolute local curvature of the mesh. b) We use geometric eigenmodes derived from the five surfaces in panel a to reconstruct each of the seven task contrasts; text colors of the labels in panel a correspond to the plots’ line colors in panel b. c) Schematic of workflow to randomly generate 100 bulbous shapes, compute 200 geometric eigenmodes for each shape, and reconstruct the seven unperturbed task contrast maps. d) Results of the data reconstruction using the bulbous shape geometric eigenmodes, following the same layout as in Figure 1, panel b.

The authors’ model of cortical wave propagation (e.g., Fig. 4 in Pang et al.) does consider long-rage axonal projections, albeit via the inclusion of isotropic distance-dependent connectivity (see equations S6-S9 in their Supplementary Material). Although structural connection weights decay exponentially with distance (Roberts et al., 2016), a simple proximity rule fails to account for the marked heterogeneity and specificity of macroscale white matter connectivity (Markov et al. 2013; Betzel & Bassett 2018).

We recognize that an *understanding* of spatially patterned brain activity will be centered on different objectives, like the reconstruction of data or a model’s corroboration of neuroanatomy, for example. We look forward to research that brings these objectives further into alignment. Such work could focus on integrating geometric constraints with the detailed topography of macroscale white matter tracts (Jbabdi et al. 2015). Such a synergistic approach might better reconcile the model’s accuracy in reconstructing observed phenomena, such as the segregation of the dorsal and ventral processing streams in Fig. 4, with the underlying reality of precise and heterogeneous anatomical connectivity (Passingham et al., 2002).

A second concern relates to the specificity of the basis sets for explaining brain function. Imagine that we accept the results of Pang et al. as conveyed in the manuscript—that excitation of geometric modes provides a more accurate and parsimonious explanation of brain function. We would expect that these modes exhibit specificity—i.e., while they should be well-suited for explaining observed patterns of brain activity, they should perform poorly in explaining randomly oriented activity patterns, uncoupled from the underlying cortical anatomy. Conversely, we might expect that modes derived from non-brain-like shapes, such as a spherical or bulbous surface, would be less accurate descriptors of brain activity than the cortical geometric modes. However, we find that neither of these expectations are met.

The modes derived by the authors are equally adept at explaining randomly rotated activation maps of brain activity (**Fig. 1**; see **Supplementary Methods** for details of these analyses). Notably, the connectome eigenmodes similarly lack specificity (**Supplementary Fig. 1**), demonstrating that high dimensional eigenmodes, geometric or otherwise, are flexible tools for modeling data generally. These observations suggest that modes derived from cortical geometry may be tuned generically to the spatial frequencies of smooth and continuous maps inherent to functional magnetic resonance imaging data but exhibit relatively little specificity to the orientation of the brain activation maps (but see also **Supplementary Fig. 3** for evidence of geometric modes’ effectiveness of reconstruction at high spatial frequencies).

Furthermore, geometric modes derived from the sphere, other brain shapes, or randomly perturbed surfaces explain brain activation maps as well as the modes used by the authors (**Fig. 2** and **Supplementary Fig. 2**). This suggests that the actual shape of the brain, including the contours of its folding patterns, is not necessary to parsimoniously describe brain activity maps, using the framework employed by Pang et al (see also Robinson et al. 2016, for example). The authors’ analyses (Supplementary Fig. 4 in Pang et al.) also demonstrate that individual-specific geometric eigenmodes contribute nominally to differences in reconstruction accuracy. Collectively, these observations suggest that the modeling approach may be insensitive to the specific shape of the cortex, and likely more adapted to the spatial adjacency or smoothness of the data.

In conclusion, Pang et al. (2023) present an interesting framework for representing brain function based on well-established physical models and with clear applications for neuroscience. The remarkable reconstruction accuracy of the geometric eigenmodes, using only a fraction of the available dimensionality, demonstrates that these methods can describe patterns of fMRI data in a compact manner. We do not doubt that this approach, and similar frameworks (Atasoy et al. 2016; Cabral et al. 2023; Luppi et al. 2023), can provide insight into spatio-temporal brain dynamics, particularly as they relate to the tradeoffs shaping brain evolution. However, we are concerned how the model of brain dynamics put forth in Pang et al. inadvertently overlooks topologically complex and long-distance white matter connectivity, beyond what can be captured by a non-specific exponential distance rule. Our critique extends to the apparent flexibility of the methodology, which as we show, does not seem to depend uniquely on the orientation and shape of the brain.

## Disclaimer

The presented opinions are those solely of the authors and do not necessarily represent the opinions of the National Institutes of Health.

## Acknowledgements

J.F., D.A.H., J.G-C, and P.A.B were supported by the Intramural Research Program of the National Institute of Mental Health (annual report ZIAMH002783). This research was supported in part by Lilly Endowment, Inc., through its support for the Indiana University Pervasive Technology Institute.

## Supplementary Methods

### Spin-test null modeling

We downloaded 200 geometric eigenmodes employed in Pang et. al, which were derived from the triangular surface mesh representation of the midthickness human cortical surface (left hemisphere). We additionally downloaded the seven key task activation contrast maps projected to the cortical surface as depicted in Figure 1e of Pang et. al, which were processed and openly shared by the Human Connectome Project (Elam, 2021). For each of the seven maps, data was randomly rotated in a spherical manner (5000 iterations) using the BrainSpace toolbox (Vos de Wael, 2020). This method of null modeling randomizes vertex location, yet preserves spatial structure, of the data on the cortical surface (Alexander-Bloch, 2018; Fig. 1a). The unperturbed geometric eigenmodes were used to predict the spun data (excluding vertices corresponding to the spun medial wall) with increasing numbers of eigenmode dimensionality, as performed in Pang et. al. We show in Fig. 1b that geometric eigenmodes predict randomized contrast maps similarly to the unperturbed contrast maps. We performed an analogous analysis as described here, using the connectome eigenmodes to predict the randomized contrast maps. In Supplemental Figure 1 we show that the unperturbed connectome eigenmodes predict randomized contrast maps similarly to the unperturbed contrast maps.

### Additional brain map null modeling

We applied two additional null modeling approaches for spatial maps. The first additional method uses the BrainSpace toolbox (Vos de Wael, 2020) to create randomized maps using the Moran randomization approach. This method was initialized with eigenvectors of the inverse geodesic distance (i.e., closeness) between all vertices of the 32k midthickness mesh, with no thresholding applied. Sampling was performed using the singleton procedure implemented in the BrainSpace toolbox. The second method also utilizes these eigenvectors, but only the first 5000 dimensions (sorted by descending eigenvalue). These 5000 modes were used as a basis set to fit to the empirical brain map data, using the same approach as Pang et al. when fitting the geometric modes to the empirical data. Following this fitting, 2500 randomly selected coefficients were sign flipped and then multiplied by the 5000-mode basis set to create surrogate data. For each method 5000 iterations were performed. Results from the additional null modeling exercises show that the low-frequency geometric modes predict random data as well as empirical data, whereas geometric modes’ reconstruction accuracy outperforms null data reconstruction after including higher frequency modes (Supplementary Figure 3). All three null modeling methods produced surrogate data that is arbitrarily correlated with the empirical data, with correlation distributions centered at zero, with ranges approximately from −0.5 to 0.5 (Supplemental Figure 4).

### Alternative brain shape eigenmodes

The midthickness human cortical surface captures the geometry of an average brain by rendering the cortical conformations in the space between the white and pial surfaces (Glasser 2013). This surface is one of several widely distributed 32k template meshes from the WashU-Minn Human Connectome Project commonly used for surface-based fMRI processing and visualization. Other commonly used surface templates with 32k vertices include the white matter, pial, very inflated, and spherical meshes, which are depicted in Fig. 2a—colored according to the absolute local curative (Cohen-Steiner & Morvan, 2003) to highlight subtle differences between their shapes. While each surface represents the brain with varying levels of smoothness and geometric contouring, the topology for all surfaces is equivalent. This means that the neighborhood relationships between vertices remain constant across these surfaces. Here, we computed 200 geometric eigenmodes from these alternative surfaces using LaPy (Wachinger, 2015; Reuter, 2006) version 0.4.1. Following Pang et al., we excluded the medial wall vertices from the geometric eigenmode construction. This operation can be performed equivalently for all shapes–even though the sphere, for example, does not technically have a medial wall–because the topology of all meshes is the same. We then subsequently used the alternative geometric eigenmodes to predict the seven key task activation contrast maps. As shown in Fig. 2b, the performance of eigenmodes derived from alternative surfaces, including the sphere, perform similarly to the midthickness surface. We also used the midthickness and spherical geometric eigenmodes to predict 10,000 contrast maps (as shown in Pang et al. Fig. 3) from the NeuroVault database (Supplemental Fig. 2)

### Arbitrarily shaped eigenmodes

We generated random bulbous shape meshes, to see if their associated geometric eigenmodes could similarly reconstruct the seven key task activation contrast maps. We started with the 32k midthickness mesh and computed Moran eigenvectors of this mesh (Vos de Wael, 2020). Briefly, this involved calculating the inverse geodesic distance (i.e., closeness) between all vertices, retraining only values with distances less than the 20% distance percentile. We then computed an eigen-decomposition on the thresholded closeness matrix to recover 10 modes of variation. A random selection (with replacement) of the modes normalized between −1 and 1 were used to modulate the x, y, and z coordinates of the spherical mesh (times a constant of 15% of the x, y, or z coordinate range), to render a new shape with wavy contours (Fig. 2c). This process was repeated 100 times and 200 geometric eigenmodes were computed for each of the shapes using LaPy (excluding the medial wall vertices as described previously). These modes were then used to predict the seven key task activation contrast maps, as shown in Fig. 2d.

**Supplemental Figure 1.**
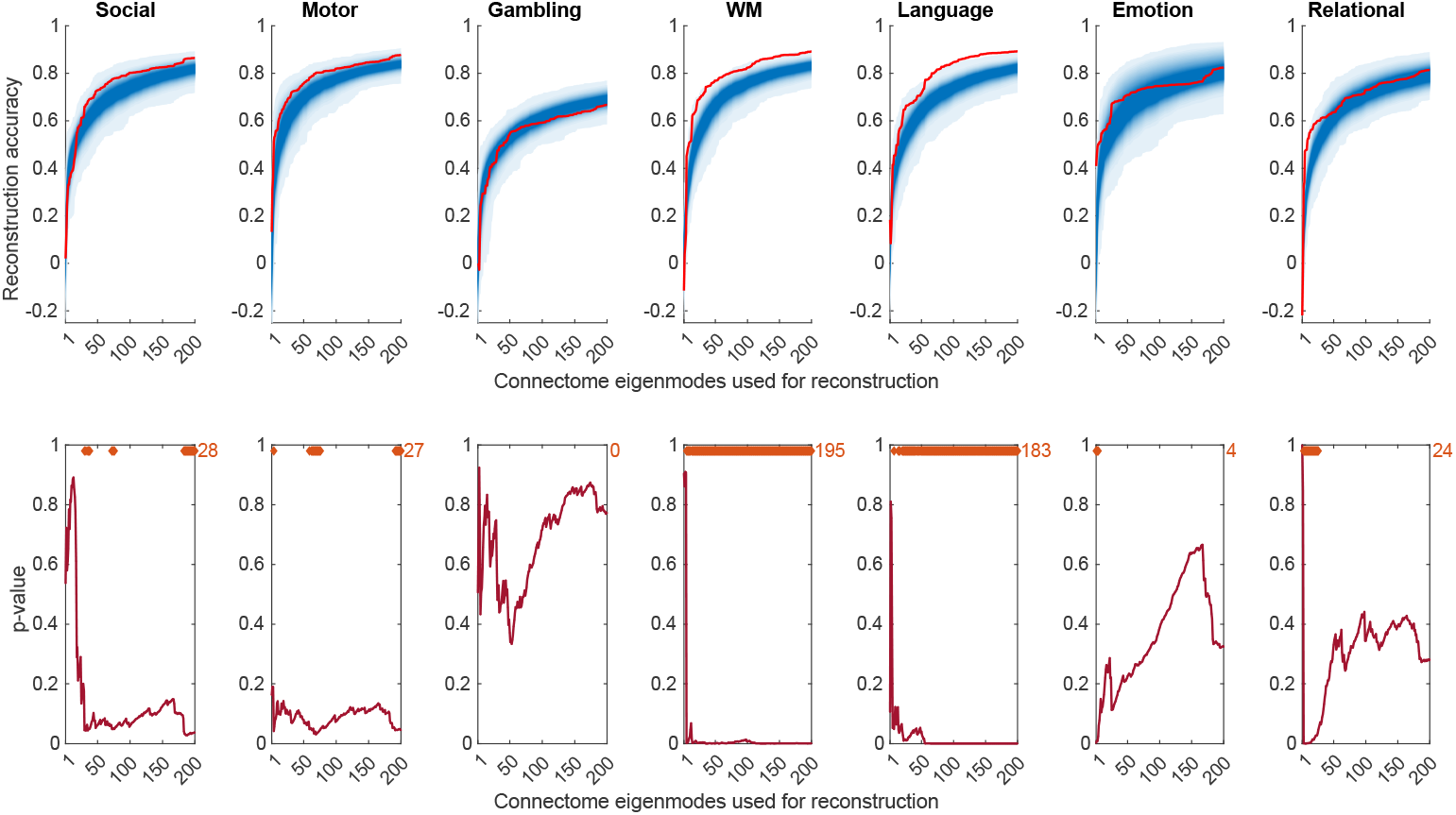
Reconstruction accuracy of connectome eigenmodes. (a) Results of the data reconstruction using the connectome eigenmodes on spun empirical data, following the same layout as in Figure 1, panel b.

**Supplemental Figure 2.**
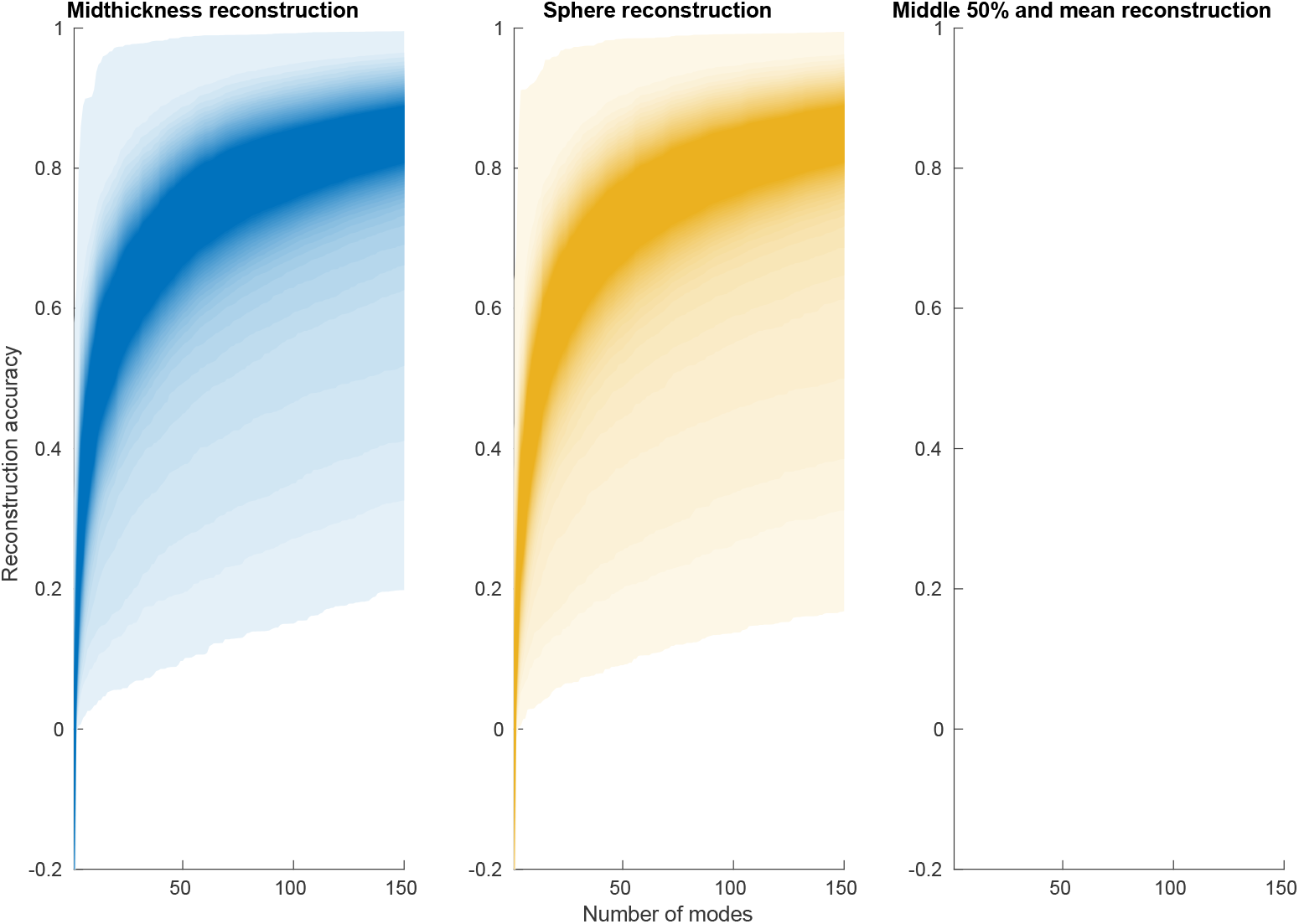
Reconstruction accuracy of 10,000 contrast maps from the NeuroVault database. We visualize the reconstruction accuracy using the geometric eigenmodes (up to 150 modes) of the midthickness (blue) or spherical (yellow) mesh to reconstruct 10,000 NeuroVault contrast maps downloaded from the repository provided by Pang et al., where the plots are shaded according to the density of the data; the far-right plot visualizes the middle 50% of distribution each overlaid, with mean values represented by thick lines.

**Supplemental Figure 3.**
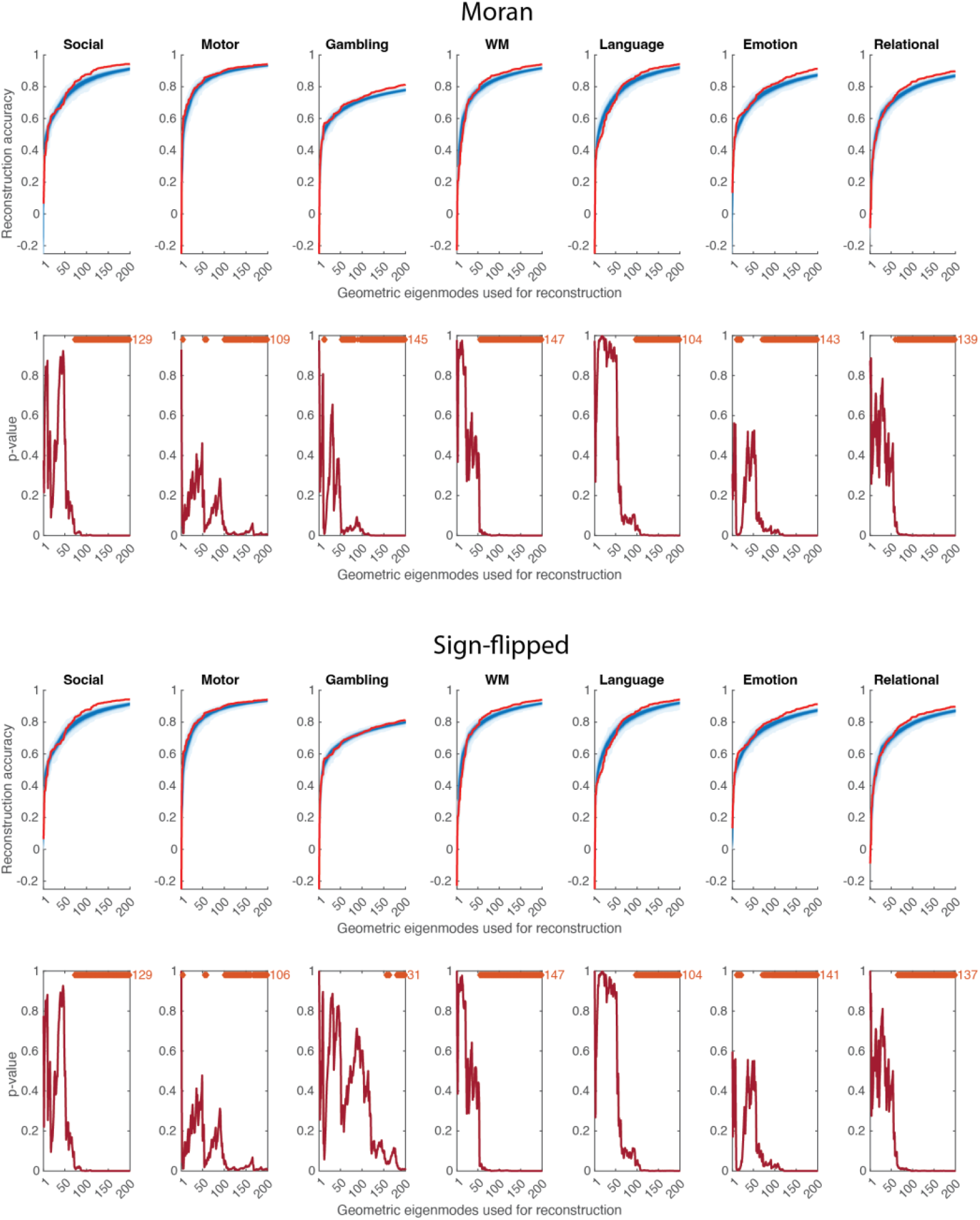
Reconstruction accuracy of geometric eigenmodes for Moran and sign-flipped surrogate data. (a) Results of the data reconstruction using the Moran (top) and sign-flipped (bottom) surrogate data, following the same layout as in Figure 1, panel b. These results show that geometric eigenmodes predict empirical data data as accurately as randomized data when using a low number of modes (up to approximately 50-100 modes), whereas geometric eigenmodes reconstruct empirical data more accurately when incorporating more modes. These results demonstrate the effectiveness of geometric modes to capture low spatial frequency signals (empirical or randomized), and further suggest that geometric modes can capture empirical structure at higher spatial frequencies more effectively.

**Supplemental Figure 4.**
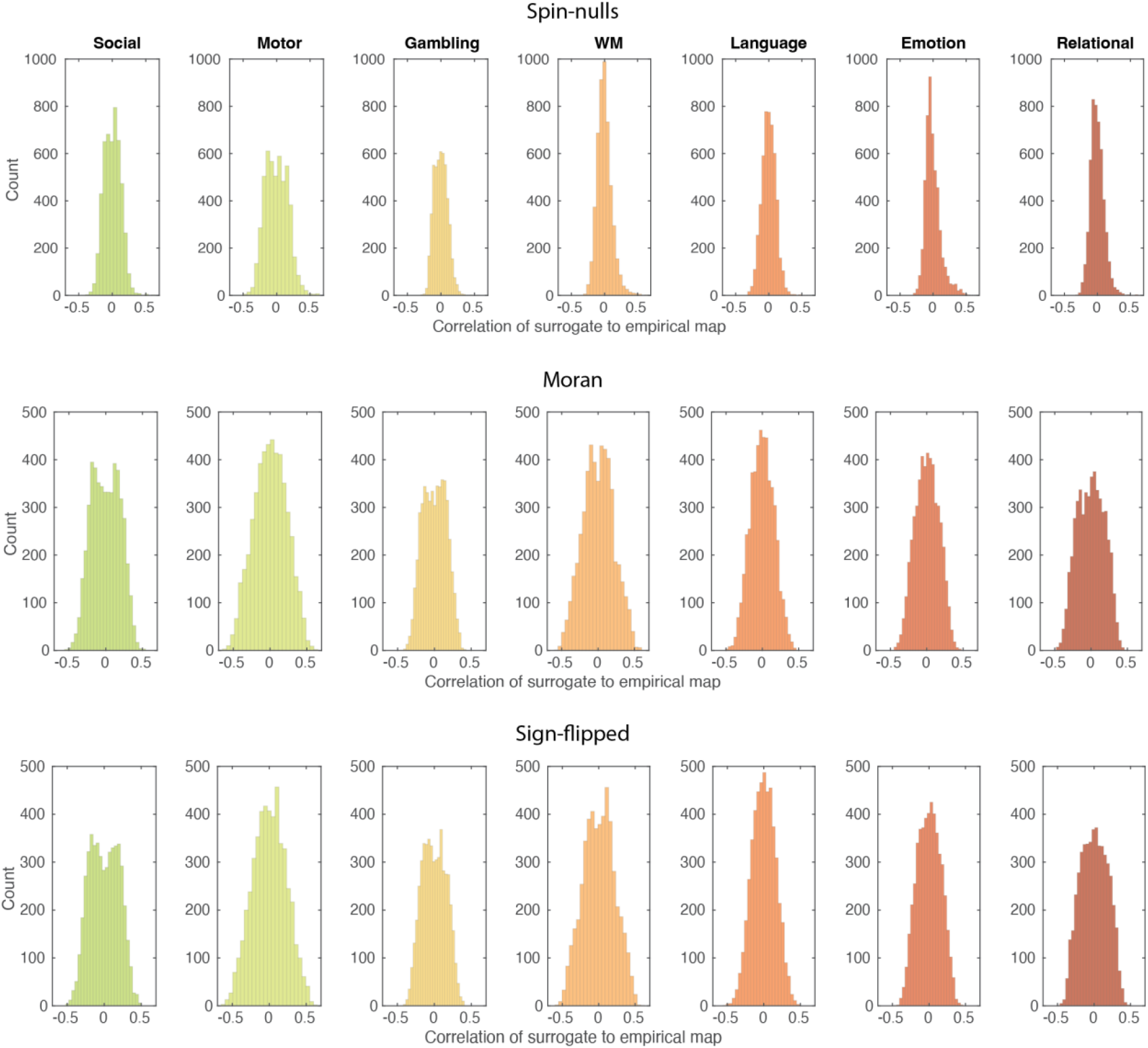
Correlation of surrogate data to empirical data. Product moment correlation of empirical data to 5000 instances of surrogate data, for each map, for each null modeling method. Note that for the spin nulls method, correlation excluded datapoints corresponding to the new placement of the medial wall.

